# Mechanical Work Accounts for Most of the Energetic Cost in Human Running

**DOI:** 10.1101/2020.09.22.309161

**Authors:** RC Riddick, AD Kuo

## Abstract

The metabolic cost of human running is challenging to explain, in part because direct measurements of muscles are limited in availability. Active muscle work costs substantial energy, but series elastic tissues such as tendon may also perform work while muscles contract isometrically at a lower cost. While it is unclear to what extent muscle vs. series elastic work occurs, there are indirect data that can help resolve their relative contributions to the cost of running. We therefore developed a simple cost estimate for muscle work in humans running (N = 8) at moderate speeds based on measured joint energetics. We found that even if 50% of the work observed at the joints is performed passively, active muscle work still accounts for 76% of the net energetic cost. Up to 24% of this cost due is required to compensate for dissipation from soft tissue deformations. The cost of active work may be further adjusted based on assumptions of multi-articular energy transfer and passive elasticity, but even the most conservative assumptions yield active work costs of at least 60%. Passive elasticity can greatly reduce the active work of running, but muscle work still explains most of the overall energetic cost.

## Introduction

It is challenging to explain the metabolic cost of human running, in part because the work and forces of the muscles are largely unknown. There is little energy dissipated by the environment, and so almost all of the action occurs within a periodic stride, with equal amounts of positive and negative work by muscles, at substantial levels of force and therefore energy cost. Although none of this information is directly measurable, there is nevertheless nearly a century of evidence about important factors such as the fundamental energetic cost of work performed by muscle, elastic energy return by tendon, and multi-joint energy transfer by muscle. These factors could potentially be combined to synthesize a plausible estimate for how much work muscles perform. This might in turn explain a substantial fraction of the overall energetic cost of running.

A first step is to quantify the mechanical work of the body. Muscles expend positive metabolic energy to perform positive and negative work, with efficiencies of about 25% and −120%, respectively (e.g., *ex vivo*^1^, for pedaling ^2^, and for running up or down steep slopes^3^ where work is largely performed against gravity). The cost of positive work is also supported by the biochemical cost of producing and using ATP for muscle crossbridges to perform work, with a net efficiency (in aerobic conditions, excluding resting metabolism) in the muscles of various animals at about 25%^4^. However, during steady, level human running, work is not readily measurable at the muscles, but rather at the body joints, as with the “inverse dynamics” technique (e.g., ^5^). Joint work does not account for multi-articular muscles, which can appear to perform positive work at one joint and negative work at another, yet actually perform no work^6–8^. The estimation of muscle work from joint work therefore depends on the assumed degree of multi-articular energy transfer^6^. Nonetheless, joint work may yet have utility, if only to place rough bounds on the work likely performed by muscle.

A second issue is elastic energy return. Muscles act in series with elastic tendons, which along with other tissues such as the plantar fascia, can store and return energy passively^9–11^. With some of the work performed on the body due to passive elasticity, running can appear to have unrealistically high positive work efficiencies of 40%^12–14^. In vivo measurements of elastic contributions in the gastrocnemius of a turkey^15^ suggest that tendon could account for about 60% of the observed joint work. But the contribution of elastic tissues to human running has been estimated in only a few cases^16–18^, and is otherwise unknown.

Elastic energy return has led to alternative measures that correlate with energy cost. For example, Kram & Taylor^19^ proposed that the cost of running is largely proportional to body weight divided by ground contact time. Referred to here as the KT cost, it presumes that much of the work observed at joints is performed passively by elastic tendon, with muscle largely acting isometrically and at high cost^20,21^. Indeed, the KT cost correlates well with metabolic cost for a variety of animals at different scales^19^, albeit with differing proportionalities for each case. But its proposed independence from work is also problematic. For example, the KT cost cannot explain the cost of running on an incline, where net work is certainly performed against gravity^3^. Even on level ground, in vivo measurements reveal muscles that do not act isometrically, but perform substantial work^16,17,22^. In addition, soft tissue deformations during running may dissipate substantial mechanical energy^23^, which can only be restored through active muscle work. Thus, work by muscle fascicles is likely still relevant to the overall energetic cost of human running.

The present study therefore re-evaluates the contribution of muscle work to running (Fig. 1). We experimentally measure the work of running, and use data from the literature to inform assumptions regarding multi-articular energy transfer, elastic energy return, muscle efficiency, and dissipative actions of soft tissues. Recognizing that the assumptions are inexact, our goal is to determine reasonable bounds, rather than an exact estimate, for the cost of work. We then test the degree to which mechanical work can explain the overall energetic cost of running.

**Fig. 1.**
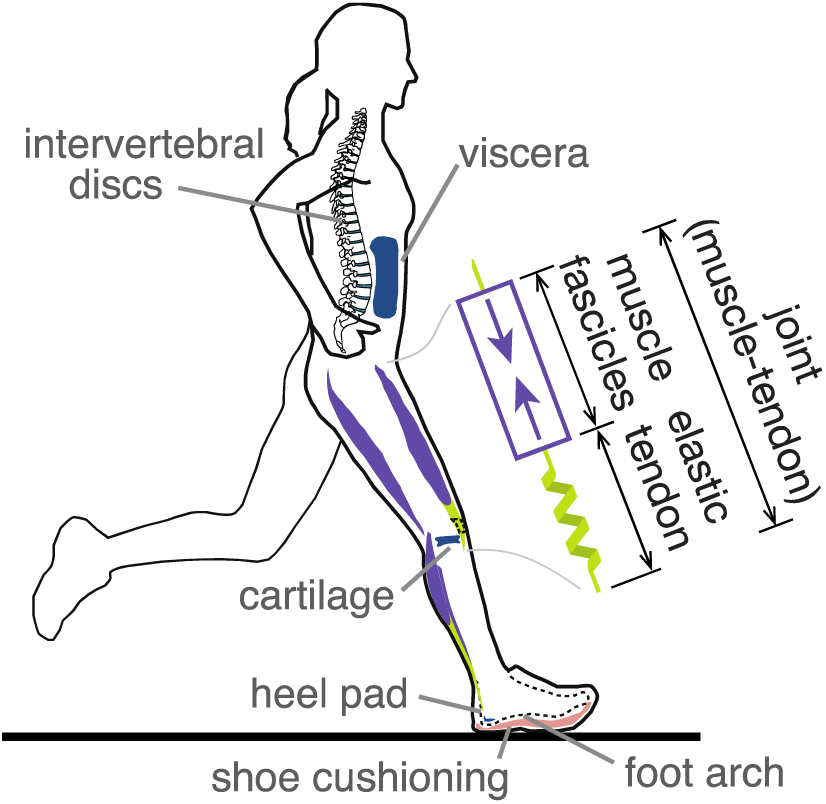
Mechanical work during human running. Muscle fascicles perform active work in series with passive elastic tendon, and the two together perform work about joints. Soft tissues such as the heel pad and the viscera also deform and dissipate energy over a stride. Traditional inverse dynamics methods quantify resultant joint work by muscle-tendon, and do not identify passive contributions from series elasticity and deformable soft tissues, nor from multi-articular muscles spanning more than one joint.

## Methods

We estimated the active mechanical work performed by the body during running, and its potential contribution to metabolic cost. We started with joint work measures using standard procedures, supplemented it with recent measures of soft tissue dissipation, and then applied simple estimates of multi-articular energy transfer and elastic energy return. Measurement were performed on healthy adult subjects (N = 8, 7 male, 1 female; 20-34 yrs) who ran at seven speeds according to each person’s comfort, in randomized order, ranging 2.2 – 4.6 m/s. Body mass M was 74.9 ± 13.0 kg (mean ± s.d.), and leg length L was 0.94 ± 0.044 m. This study was approved by the University of Michigan Institutional Review Board and all subjects gave informed consent prior to their participation.

Kinematics and ground reaction forces were recorded on a split-belt instrumented treadmill at the University of Michigan. Forces (980 Hz sampling; Bertec, Columbus, OH, USA) and motion capture (480 Hz; PhaseSpace Inc., San Leandro, CA, USA) were collected concurrently, with markers placed bilaterally on the ankle (lateral mallelous), knee (lateral epicondyle), hip (greater trochanter), shoulder (acromion of scapula), elbow (lateral epicondyle of humerus), and wrist (trapezium). Additional tracking markers were placed on the shanks, thighs, trunk, upper arm, lower arm, and upper arm, with three markers on the pelvis (sacrum, left/right anterior superior iliac spine) and two markers on each foot (calcaneus, fifth metatarsal). These data were collected for at least one minute per trial, with force data filtered at 25 Hz and marker motion at 10 Hz (low-pass Butterworth), and then applied to inverse dynamics calculations (Fig.2) using standard commercial software (Visual3D, C-Motion, Germantown, MD, USA).

These data were used to compute two kinds of mechanical work. The first was standard rigid-body joint powers, as the work per time needed to rotate and translate (via joint torque and intersegmental reaction forces, respectively) two connected segments relative to each other. We used the so-called 6-D joint power, considered robust to errors such as in joint center locations^42–44^.

The second quantity was the dissipative work performed by soft tissue deformations. Briefly, this is the difference between rigid-body joint power and the total mechanical work, defined as the rate of work performed on the COM, evaluated using ground reaction forces with no rigid-body assumptions, plus the rate of work performed to move rigid-body segments relative to the COM^23,43^. In running, this quantity is similar in magnitude to the difference between the positive and negative joint work over a stride ^23^, which itself implies that rigid body work does not capture all of the work of running.

Metabolic cost was estimated through respirometry (Oxycon; CareFusion Inc., San Diego, CA). Both O2 consumption and CO2 production were recorded on a breath by breath basis and averaged over the final three minutes of each six-minute trial, and converted to gross metabolic rate (in W). Net metabolic rate was found by subtracting each subject’s cost for standing quietly, collected before running. The subjects’ respiratory exchange ratio (RER) was measured to be 0.85 ± 0.09 across subjects, with each individual trial having an average RER of less than 1, indicating mostly aerobic conditions.

### MECHANICAL WORK AND ENERGY TRANSFER BY MUSCLE-TENDON

The work performed by joints and soft tissue deformation was used to estimate that done by the series combination of muscle and tendon. To illustrate energy transfer assumptions, we initially consider two opposing sets of assumptions—an Overestimate and an Underestimate—before introducing our intermediate measure. The Overestimate assumes no multiarticular energy transfer between joints, as if all muscles acted uniarticularly. Positive work is thus evaluated by integrating the positive intervals of each joint’s power over a stride (Fig. 2A), and then summing across all joints in the body, as if they were independent joints (IJ). Multiplying by stride frequency then yields the average rate of positive independent-joint work, 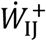. We consider this quantity to be an Overestimate because it disregards energy transfer by multi-articular muscle.

The Underestimate of work takes the opposite extreme, and assumes that simultaneous positive and negative work always cancel each other. This entails summing the powers from all the body joints at each instance in time, yielding summed joint power^43^, and then integrating the positive summed joint power over a stride. Multiplying by stride frequency yields the average rate of positive summed-bilateral (SB) joint work, 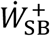 (Fig. 2B). This is considered an Underestimate of actual muscle-tendon work, because it assumes energy transfer can occur between any two joints, regardless of whether a muscle crosses those joints. The Over- and Under-estimates, 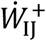 and 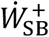, are roughly analogous to the terms “no between-segment transfer” and “total transfer between all segments” of Williams and Cavanagh^33^, except applied here to transfer between joints rather than body segments. We introduce our own intermediate muscle-tendon work estimate, termed Summed Ipsilateral (SI) work. It assumes full energy transfer across the joints on each side of the body, but not between the two sides. This is because there are no muscles that cross the legs and could transfer negative work from one leg into positive work at the other. The average rate of work 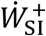 entails summing the joint powers on one side of the body at each point in time, integrating the positive intervals of this power (Fig. 2B), and then multiplying by step frequency. Of course, further examination of musculoskeletal geometry, neural activation patterns, and loading conditions could yield more intricate estimates of muscle-tendon work. But without full knowledge of individual muscle forces and displacements, we use the Summed Ipsilateral estimate as a simple and not unreasonable set of assumptions, between the aforementioned extremes.

**Fig. 2.**
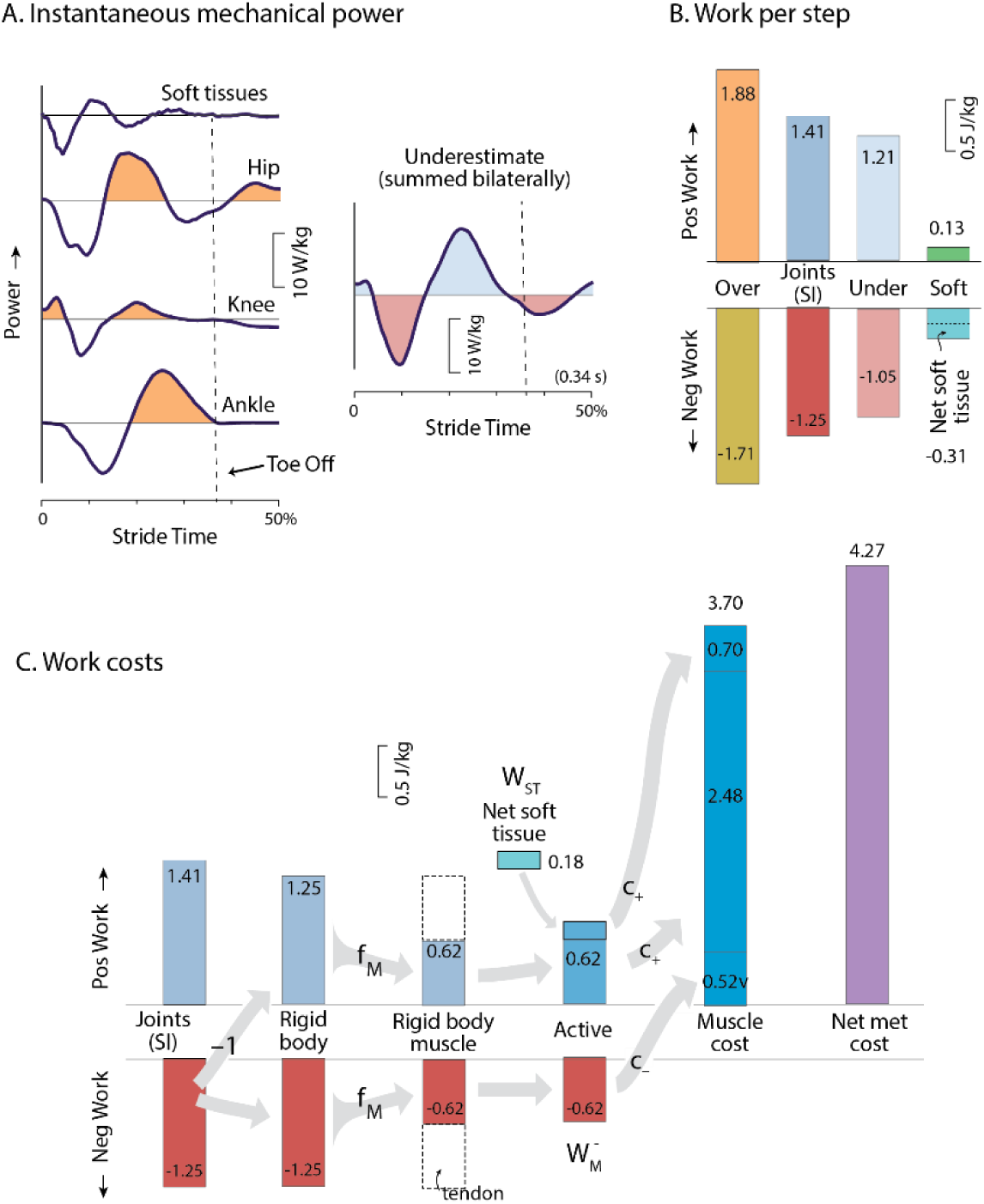
Mechanical work contributions to metabolic energy expenditure, for a representative subject (3.10 m/s). **(A)** Instantaneous mechanical power of the joints (ankle, knee, hip and upper body), and from soft tissue deformations, over one-half running stride (beginning with heelstrike). Also shown is the summation of all joint powers from both sides of the body, which is an underestimate of power **(B)** Four summary measures of work per step: Overestimate, Estimate (Summed Ipsilateral work), Underestimate, and Soft Tissue work. Positive (negative) work refers to integrated intervals of positive (negative) power. Soft tissues perform net negative (dissipative) work. **(C)** Work costs illustrate metabolic cost contributions. The magnitude of Summed Ipsilateral negative work is treated as an estimate of the joint positive and negative work performed on rigid body segments. This is multiplied by muscle work fraction f_M (provisionally 0.5) to yield work due to muscle. Active muscle work also includes positive work to offset net soft tissue dissipation. These are multiplied by costs for positive and negative muscle work (c_+ and c_-) to estimate the energetic cost due to active muscle (“muscle cost”), in this example about 86% of the net metabolic cost of running.

### METABOLIC COST OF MUSCLE WORK

We define two quantitative parameters to link muscle-tendon mechanical work to energy expenditure. The first is the proportion of work performed actively by muscle vs. passively by tendon, and second is the metabolic cost at which the active work is performed. The proportion is defined as *f*_*m*_, the fraction (ranging 0-1) of muscle-tendon work performed by muscle fascicles, such that

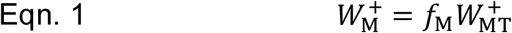

Where 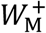 is the positive work of muscle fascicles and 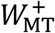 is the positive work of muscle-tendon (applying the proposed Summed Ipsilateral measure, or the Over- or Under-estimate assumptions), and analogously for negative work. In vivo measurements suggest a variety of possible values for *f*_*m*_, for example 0.40 for turkey gastrocnemius ^15^, and 0.26-0.56 for two muscles of running dogs^45^. For humans, cadaver data suggest 0.52 for the Achilles tendon and foot arch^10^. Other indirect data suggest a range of 0.4 – 0.625 (Cavagna et al., 1964; Cavagna & Kaneko, 1977), depending on energy transfer assumptions. The correct value is unknown, and almost certainly varies with muscle group, loading conditions, and speed. We use a single parameter *f*_*m*_ to summarize an overall effect for all muscles, and adopt a provisional value of 0.5, while allowing for other possible values.

We characterize the metabolic cost of muscle work with separate parameters for positive and negative work. The positive work cost *c*_*+*_ is defined as the metabolic energy cost of producing a unit of active positive work, equivalent to the inverse efficiency of pure positive work. An analogous cost *c*_−_ is defined for the metabolic cost of negative work. We adopt provisional values for *c*_*+*_ and *c*_ of 4.00 and −0.83, respectively, equivalent to efficiencies of 25% and −120% (e.g., ^46^).

The overall energetic cost of this work *E*_*work*_ is summed for rigid body and soft tissue contributions (graphically depicted in Figure 2C). Soft tissues dissipate net energy (yielding negative 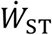), and muscles must actively perform net positive work to compensate for those losses. The positive cost of making up for such dissipation is therefore *c*_*+*_|*W*_*ST*_|. The cost of rigid body work is estimated from the magnitude of negative work from inverse dynamics 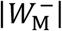, multiplied by the costs for both positive and negative work. These summed contributions yield

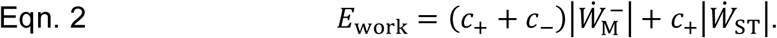

This energetic cost per stride is then multiplied by stride frequency to yield metabolic power *Ė*_work_ due to active work.

To account for differences in subject size, data were non-dimensionalized using body mass *M* leg length *L*, and gravitational acceleration *g* as base variables. Mean power and work normalization constants were *Mg*^3/2^*L*^1/2^ = 2184 W and *MgL* = 678 J, respectively. The mean running speed normalization constant was *g*^1/2^*L*^1/2^ = 3.04 m/s. All averaging and statistical tests were performed with dimensionless quantities. In figures, data were plotted with dimensional scales in SI units, using the mean normalization constants.

Statistical tests were performed as follows. We used a linear least-squares fit to relate running speed to mechanical or metabolic rates, and then used Eqn. 2 to estimate the metabolic cost attributable to work. We also used linear regression to test how other work measures and the KT cost are related to metabolic rate. All regressions were performed allowing each subject an individual constant offset, while constraining them all to a single linear coefficient. Significance of work trends across running speeds was tested with repeated measures analysis of variance (ANOVA), and differences between work measures with repeated measures t-tests (α = 0.05).

## Results

We found that mechanical work rates and metabolic rate all exhibited typical and fairly linear increases with running speed. In terms of standard joint powers (Fig. 2A, representative data), the ankle, knee, and hip powers far exceeded that for the upper body. Soft tissues produced power similar to a damped oscillation (reported previously;^23^), and the Over- and Under-estimates of power bracketed the intermediate estimate, as expected. This was also true for the overall Over- and Under-estimates of positive and negative work per stride (Fig. 2B); soft tissues produced net negative work. These observations were consistent across subjects and running speeds (Fig. 3). As expected, the proposed Summed Ipsilateral work rate increased with running speed (Fig. 3A), and was between the expected Overestimate and Underestimate. Net soft tissue work rates were negative and increased in magnitude with speed.

**Figure 3.**
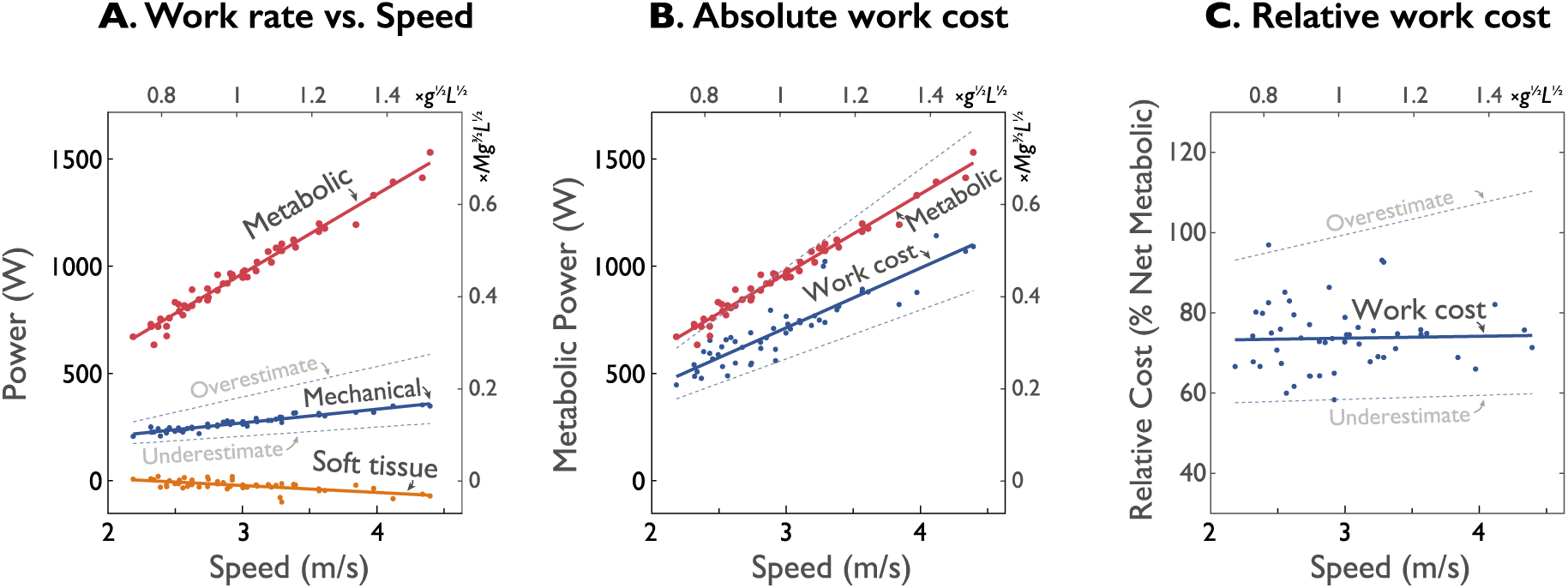
Mechanical work and estimates of absolute and relative metabolic cost vs. speed (*N* = 8). (**A)** Average positive work rates: Mechanical work (using Summed Ipsilateral estimate), net Metabolic rate, and net Soft tissue work rate. Also shown are Over- and Under-estimates of work (dashed gray lines) assuming no work transferred between joints by multiarticular muscles, and full transfer, respectively. (**B)** Estimated metabolic cost of mechanical work, based on each work rate, along with soft tissue deformations, muscle work fraction, and muscle work cost. (**C)** Relative metabolic cost of mechanical work, showing each cost as a fraction of net metabolic rate. Axes shown include dimensional units, as well as dimensionless units (top and right-hand axes) using body mass, leg length, and gravitational acceleration as base units.

The estimated metabolic cost for performing that work was substantial. Applying elastic contributions, the metabolic cost for performing active work (Eqns. 1 & 2) ranged about 500-1000 W over the speeds examined, compared to an overall net metabolic rate of 700 – 1500 W (Fig. 3B). In relative terms (Fig. 2C), work accounted for about 76% of net metabolic rate (Fig. 3C), with little dependence on running speed (slope = 0.10 % per 1 m/s change in speed). In contrast, the Overestimate of work yielded a much higher proportion (slope = 7.1 % per 1 m/s change in speed), actually exceeding 100% of net metabolic rate at most speeds. The Underestimate yielded a fairly constant proportions of about 61% (slope = 0.62 % per 1 m/s change in speed).

To facilitate evaluation of assumptions, the fraction of metabolic cost explained by work is illustrated as a function of parameters (Fig. 4). Here we use an overall cost for combined positive and negative work, *c*_±_ = *c*_*+*_ − *c*_−_, with nominal value 4.83. This is nominally paired with muscle work fraction *f*_*m*_ of 50%. With these values, the proportion of metabolic cost explained by work was 61% for the Underestimate, 76% for Summed Ipsilateral, and 106% for Overestimate, respectively, across the observed running speeds. Here we examine two extremes for alternative assumptions. One is to assume a considerably lower fraction of muscle work, *f*_*m*_ = 0.38, which would yield a lower fraction of metabolic cost explained, of 43%. On the other hand, assuming that muscle performs more work, *f*_*m*_ = 0.65, yields an unrealistic explained fraction of 1.53.

**Figure 1.**
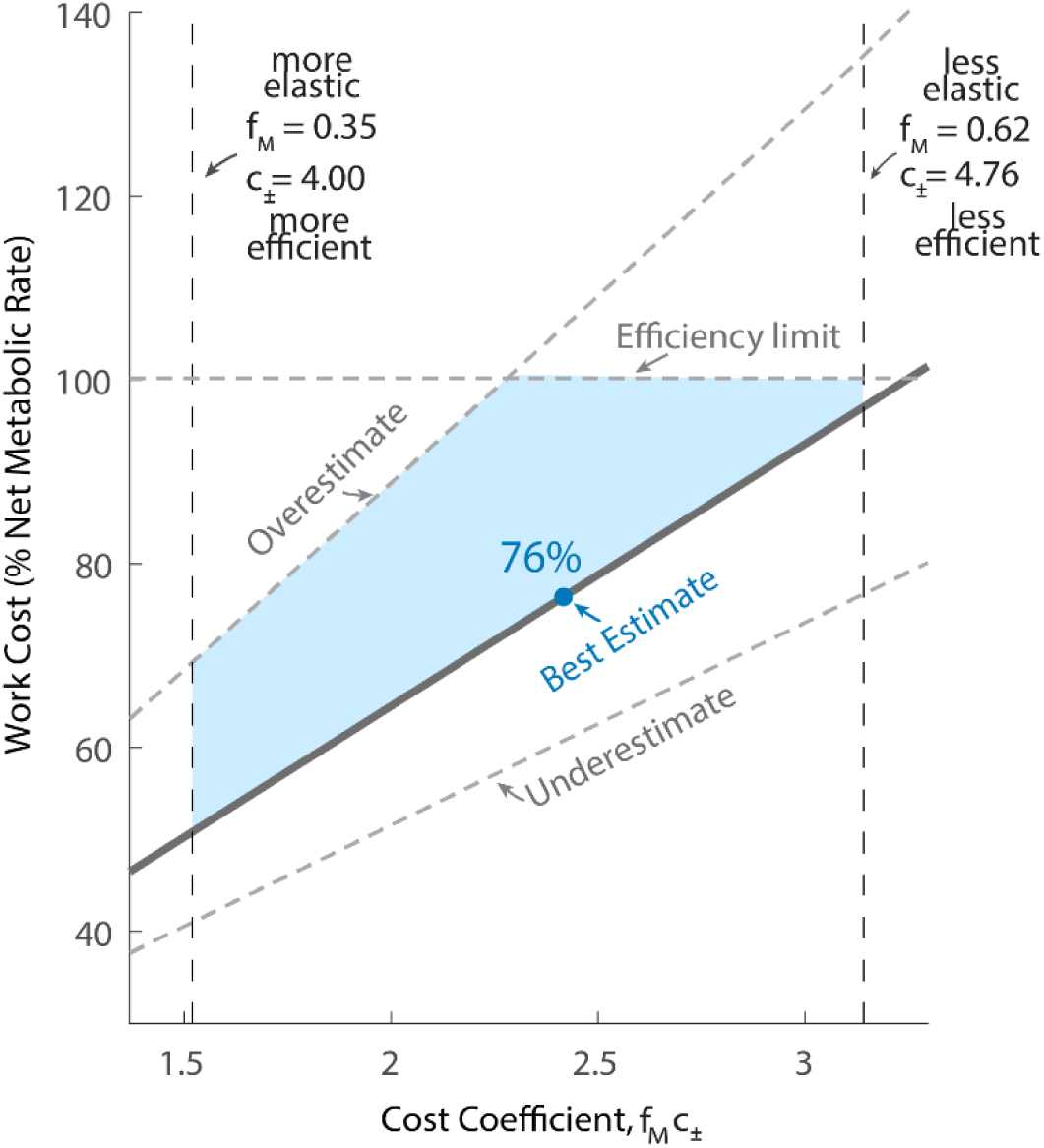
Work cost as a function of cost coefficient for running at 3 m / s. Relative work cost is estimated metabolic cost of mechanical work divided by overall net metabolic cost. Cost coefficient is defined as fraction of work attributable to muscle from overall muscle-tendon work, multiplied by cost of active work *c*_±_. Boundaries are shown for extreme assumptions. Overestimate is for Independent Joints assumption, where muscles only act uniarticularly; underestimate is for Summed Bilateral joint assumption, where work can be transferred from one side of the body to the other. Left and right boundaries are for extremes in muscle work fraction, 35% and 62%, respectively, with constant cost of work. The proposed work estimate (Summed Ipsilateral joints), along with a muscle fraction of 50%, yields 76% of the metabolic cost of running is attributable to active work by muscle.

Using the nominal efficiency of *c*_±_ along with the Summed Ipsilateral cost for work, active work to compensate for soft tissue dissipation accounted for an increasingly larger proportion of the metabolic cost due to work. At a nominal speed of 3 m/s, soft tissue compensation increased the estimated metabolic cost due to work by 23.3%, from 3.00 J/kg to 3.70 J/kg. Whereas at the highest speed of 4.6 m/s, soft tissue compensation increased the estimate of cost due to work by 31.5%, from 3.82 J/kg to 5.03 J/kg.

## Discussion

We had sought to re-evaluate the degree to which mechanical work performed by muscle can explained the net metabolic cost of running. We considered three sets of assumptions to translate joint work estimates into metabolic cost: how energy is transferred between joints by muscle, how much work is performed passively by tendon, and how much metabolic energy is expended to perform muscle work. Using nominal assumptions derived from the literature, we found that about 76% of the metabolic cost of running is attributable to muscle work. We next discuss how our estimates may be interpreted, and how they could be affected by alternate assumptions.

One contributor to the high work cost is dissipation by soft tissues. The dissipation is not typically measured in inverse dynamics analysis, nor incorporated into estimates of metabolic cost. In a typical inverse dynamics analysis, the only work is performed about joints acting between rigid segments, leading to an imbalance of work^23,24^, with more positive than negative work. In fact, soft tissue deformation largely explains the joint work imbalance^23^. For example (representative subject, Figure 2), soft tissues dissipated 0.18 J/kg, explaining much of the positive/negative work discrepancy of 0.16 J/kg at 3.1 m/s. Active work to make up for this dissipation accounted for 0.7 J/kg (16%) of the entire 4.27 J/kg of the net metabolic rate. And at faster speed of 4.6 m/s, that fraction increases to about 31%. Faster speeds entail higher impact between leg and ground, and more energy dissipation. The work to compensate for soft tissue energy dissipation costs substantial metabolic energy.

Another contributor is active work in tandem with passive elasticity. Series elasticity is recognized to perform substantial work passively, and thus to play an important role in running energetics. But even with passive elasticity, our results suggest that the remaining work attributable to muscle accounts for much of the overall energetic cost. This is based on an assumed muscle work fraction *f*_*m*_, provisionally set to a nominal value of 50%, for which far different values might be appropriate. For example, the plantaris and gastrocnemius of hopping wallabies have a range from only 3-8%^25^. In human, the Achilles tendon appears to facilitate a low muscle work fraction^14,22^. However, many other muscles also participate in running, not all under conditions ideal for tendon elastic work. It is therefore helpful to use the parameter study (Fig. 4) to evaluate other candidate assumptions.

Another factor in our energy estimate is the energetic cost of muscle work. This is mainly for positive work, and attributable to crossbridge cycling^26^. Thermodynamic principles dictate that this cost likely exceeds *c*_*+*_ = 4 (or efficiency does not exceed 25%), due to the biochemical costs of ATP production and for crossbridge work^27^. We did not include other effects such as muscle co-contraction, isometric force production, or calcium pumping^28^, which would generally be expected to cost energy, and could be lumped into the remaining fraction of energy cost (24%) not explained by fascicle work. We also assumed a small but positive energetic cost to negative work. An extreme assumption would be zero cost for negative work, which would reduce the estimated metabolic cost for work from 76% to about 63%, still a majority of overall metabolic cost.

We also examined alternative assumptions for energy transfer by multi-articular muscles. Although generally unknown in humans, measurement of muscle forces in cat locomotion show significant energy transfer from the ankle to the knee during collision, and from the knee to the ankle during push-off^29^. We therefore consider it unrealistic to assume no such transfer in humans, hence the label of Overestimate for the individual joints (IJ) estimate of work. Indeed the IJ estimate would yield an entirely realistic apparent mechanical efficiency of 102% for running at 3 m/s (Fig. 4). On the other hand, the Underestimate is likely too low, since it assumes that negative work at any joint could be transferred perfectly to positive work at any other joint in the body. We have therefore presented the Summed Ipsilateral (SI) assumption as a better, yet likely low, estimate for work performed by muscles. It has long been recognized that energy transfer can occur between joints of an individual leg^6,29–31^. Our own estimates could be improved with more direct muscle measurements from human.

Our findings could inform other estimates of mechanical work. Others have used independent joint work to evaluate apparent efficiency during locomotion^14,32^, for example yielding unusually high running efficiencies of 35-40%^14^, which they largely attributed to series elasticity at the ankle. But we also believe some of their observed work may be an Overestimate, due to multi-articular energy transfer. Our preferred estimate using summed ipsilateral joint work is more similar to the segmental energy transfer approach of Williams & Cavanagh^33^, except using work at joints rather than between segments, and including soft tissue work not been previously considered. This facilitates estimation of metabolic cost contributions (eqn. 2) with only two main parameters (*f*_M_ and *c*_±_) lumped into the cost coefficient. We anticipate that further measurements of muscle and tendon action *in vivo* will inform better estimates of cost contributions such as work.

There are certainly other costs for running, not attributable to work. Examples include a cost for producing force in the absence or regardless of mechanical work^19–21^, or due to the rate at which force is generated^34^. We evaluated the KT cost (Fig. 5) proportional to body weight divided by ground contact time^19^, which correlates quite well with metabolic cost. But a number of measures, including various estimates of work, also correlate well (Fig. 5). We consider it more mechanistic for a cost to depend on applied muscle force or work, rather than parameters such as body weight. For example, “Groucho running” ^on flexed knees, 35^ costs 50% more energy than normal running, whereas the KT cost would predict a decrease, due to increased ground contact time. We suspect that the high cost of Groucho running is due to greater muscle forces and work with flexed knees (e.g., ^36^) even though body weight remains unchanged. Furthermore, reinterpretation of the KT cost reveals that it could be equivalent to a cost of performing mechanical work under appropriate assumptions (see Supplementary Appendix S1). Work is certainly needed to accelerate during running, or to ascend an incline, and it appears to account for a majority of the cost for level ground. We do acknowledge other costs, potentially for isometric force production, but mainly for the 24% of energy not explained by work.

**Figure 5.**
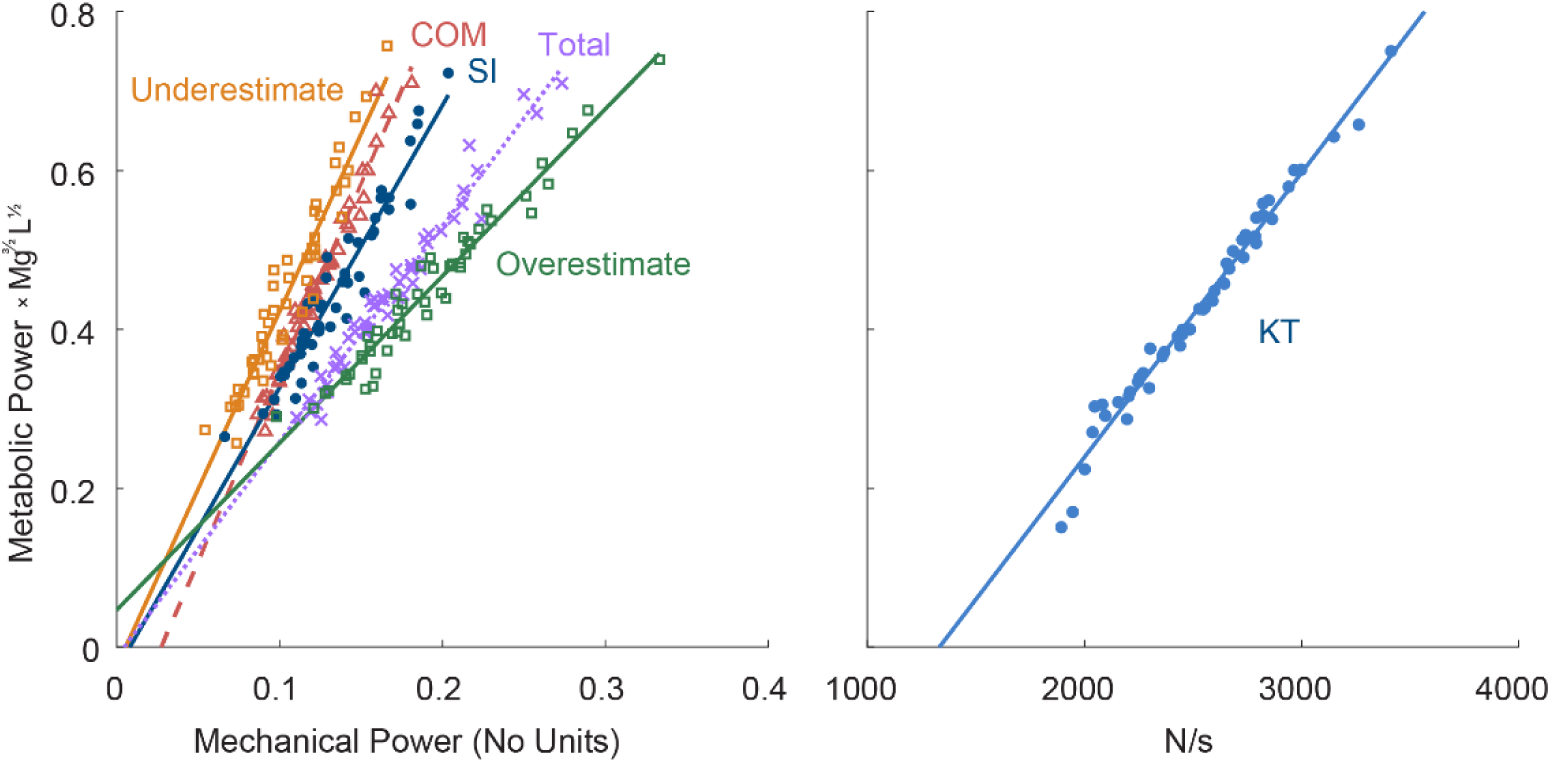
Sample correlates of metabolic cost. **A)** Correlates: Summed Ipsilateral (SI) work, positive COM work rate, Total mechanical work, Underestimate of joint work (assumes full energy transfer), and the Overestimate of joint work (assumes joint independence). **B)** The KT measure of body weight divided by ground contact time (Kram & Taylor, 1990) compared to metabolic cost. All measures correlate well (r^2^ >0.9) with metabolic cost. Power is plotted in terms of normalization units, *Mg*^3/2^*L*^1/2^.

The present study has a number of limitations. Our results are specific to humans running at a limited range of speeds, and it remains to be seen how well work can explain energy cost over a wider range of speeds. In particular, work may be less explanatory for other animals, particularly smaller ones where muscles are turned on or off more quickly^37^. Such force cycling costs may be applicable to humans as well^34,38,39^. And even though mechanical work should help to explain the cost of incline running, that remains to be tested. The present model is also limited to estimating metabolic cost based on motion data, whereas a more comprehensive and mechanistic model would include body dynamics and predict both motion and energy cost.

But the primary limitation is in the cost coefficient, which attempts to aggregate information from empirical data. Better estimates could be obtained as *in vivo* measurements of muscle state (e.g. ultrasound^40^) and series elastic energy storage (e.g., ^41^) become available. Still better would be to dispense with the cost coefficient in favor of detailed information about each individual muscle, including differences in fiber type and function. We expect improved estimates of elastic contributions, energy transfer, and the cost of performing work to lead to better explanation of the cost of running. However, based on current evidence, it appears that even though series elasticity performs a major role in running, active mechanical work still explains a majority of the metabolic cost in running.

**Table 1.** The cost coefficient represents how much metabolic energy a unit of mechanical work costs. The cost coefficient is calculated by taking into account the amount of work performed by tendon relative to muscle, and the efficiency of positive and negative muscle work. A range of cost coefficients between 1.8 and 3.2 were found by consulting experimental data from the literature. 1. (Cavagna et al., 1964) 2. (Asmussen and Bonde-Petersen, 1974) 3. (Margaria, 1968) 4. (Abbott et al., 1952)

| | Positive Work Cost<br>$c_+$ | Negative Work Cost<br>$c_-$ | Net Work Cost<br>$c_{\pm} = c_+ - c_-$ | Muscle Work fraction<br>$f_M$ | Cost Coefficient<br>$f_M c_{\pm}$ |
| --- | --- | --- | --- | --- | --- |
| Upper Bound | 4.00<sup>3,4</sup> | -0.83<sup>3,4</sup> | 4.83 | 0.65<sup>2</sup> | 3.14 |
| Lower Bound | 4.00<sup>3,4</sup> | 0 | 4.00 | 0.38<sup>1</sup> | 1.52 |

**Table 2.** Linear relationships between running measurements and metabolic cost. Shown are slope and offset from linear regression (in dimensionless units), along with *r*^2^. Each work measure is an of mechanical work performed at the joints or on the COM. KT refers to cost proposed by Kram & Taylor 1990.

| Measurement | Slope | Offset | $r^2$ |
| --- | --- | --- | --- |
| Estimate: S. Ipsilateral (SI) | 3.55 | -0.0285 | 0.92 |
| Overestimate: Indiv Joints (IJ) | 2.10 | 0.0467 | 0.96 |
| Underestimate: S. Bilateral (SB) | 4.45 | -0.0242 | 0.90 |
| COM work rate | 4.75 | -0.130 | 0.97 |
| Total mechanical | 2.71 | -0.0132 | 0.97 |
| Kram & Taylor (KT) | 3.58E-4 | -0.480 | 0.97 |

## Acknowledgements

This work supported in part by Natural Sciences and Engineering Research Council of Canada (NSERC Discovery Award, Canada Research Chair Tier 1) and Dr. Benno Nigg Research Chair.

## Author Contributions

R. R. collected the data, analyzed, and wrote the main text of the manuscript. A. D. K. conceived the experiment, guided the analysis of the data, and edited and wrote portions of the manuscript.

## Additional Information

The authors declare no competing interests.

## Supplementary Appendix S1

Here we use a dimensional analysis to show how the KT cost model^1^ based on body weight and ground contact time could be considered equivalent to a cost of performing work. In fact, it is equivalent to evaluating the present study’s cost of performing muscle work, using three assumptions: (1) constant speed locomotion, (2) running on level ground, and (3) a single value for the cost coefficient (lumping together the cost of active muscle work and series elastic contribution). The KT model states that metabolic cost is proportional to body weight *M*_*g*_ divided by ground contact time *t*_*c*_:

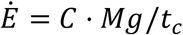

Where *C* is a constant of proportionality. Integrating this equation over stance phase results in the amount of energy used by the body during the stance phase of one step:

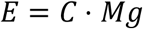

Since *E* has units of energy, *C* must have units of length. The constant also includes an implicit cost of performing the work, which converts from mechanical to metabolic energy. As we have seen, both the efficiency of muscle performing negative and positive work *c*_±_, and the proportion of work done by muscle *f*_M_ (as opposed to tendon) are factors in energy cost, and can be summarized using a cost coefficient, the product *f*_M_*c*_±_. Expanding the constant of proportionality to include these terms without a loss of generality, the constant *C* may be considered equivalent to

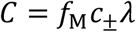

where *λ* is a parameter with units of length since the cost coefficient *f*_M_ *c*_±_ is dimensionless. Updating and rearranging the equation for the energy cost of a step results in

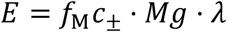

The first term *f*_M_ *c*_±_ is the cost coefficient, the second term is the average load the ground applies on the body *Mg*, and the third term *λ* is therefore the distance displaced by the load, by the definition of mechanical work. This parameter *λ* represents the distance travelled by the body’s center of mass along the direction of the applied ground force. We calculated this value for the running data used in this paper and found it to be equal to 0.090 ± 0.015 *m*. Examining the value for *C* reported for humans, it has an average value of 0.262 m across a range of running speeds from 2 to 4 m/s^2^, similar to the running speeds in our dataset. Dividing this value reported for *C* by the average distance travelled by the body *λ*, we get a cost coefficient of *f*_M_ *c*_±_ = 2.91, slightly higher than the value 2.41 we used, and within the range of reasonable values for the cost coefficient (Figure 4). This shows that the value of *C* is a single parameter that represents the effects of three distinct physical phenomena: energetic cost of work, series elasticity, and displacement of the COM along the direction of the ground force.

It therefore appears that the KT model is a special case of the cost of generating mechanical work, under steady state, level ground locomotion. To generalize this equation across terrains, accelerations and decelerations, and types of locomotion would be a matter of separately assessing the amount of distance travelled by the body when performing both negative and positive work, and taking into account the different efficiencies for both types of work. Our cost model (equation 2) provides the framework for doing so, and it applicable even when locomotion is not constant speed nor on level ground.

## Notes

### Competing Interest Statement

The authors have declared no competing interest.

